## Supplementary Figures 1-7 for "Strong pathogen competition in neonatal gut colonisation"

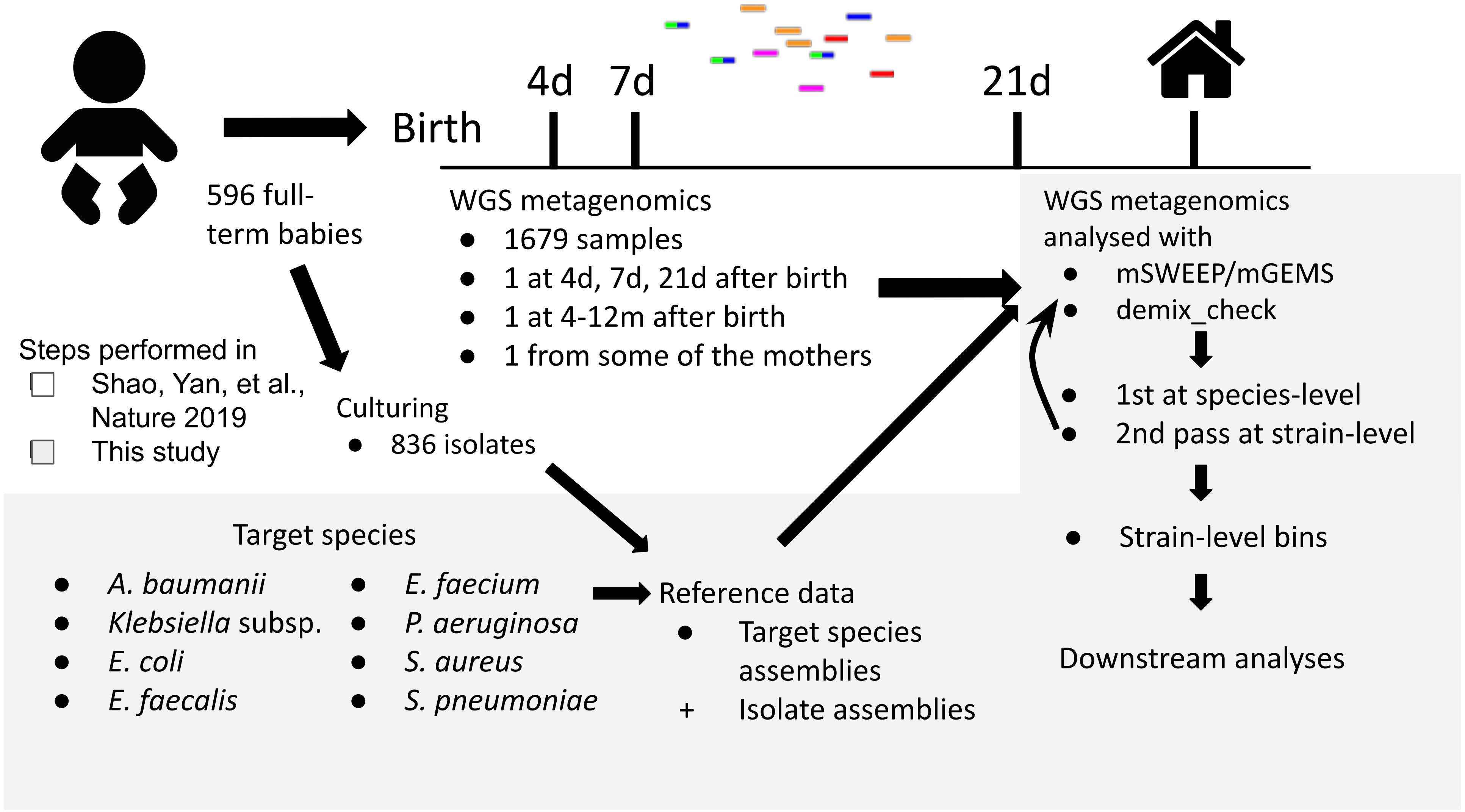


**Supplementary Figure 1 Flowchart describing the data collection and analysis steps.** The figure shows an overview of how the sequencing data was collected and analysed. Sampling and sequencing was performed in the source study (Shao, Yan, et al., Nature 2019; white background in the figure), and the analysis and reference gathering were performed in this study (grey background).

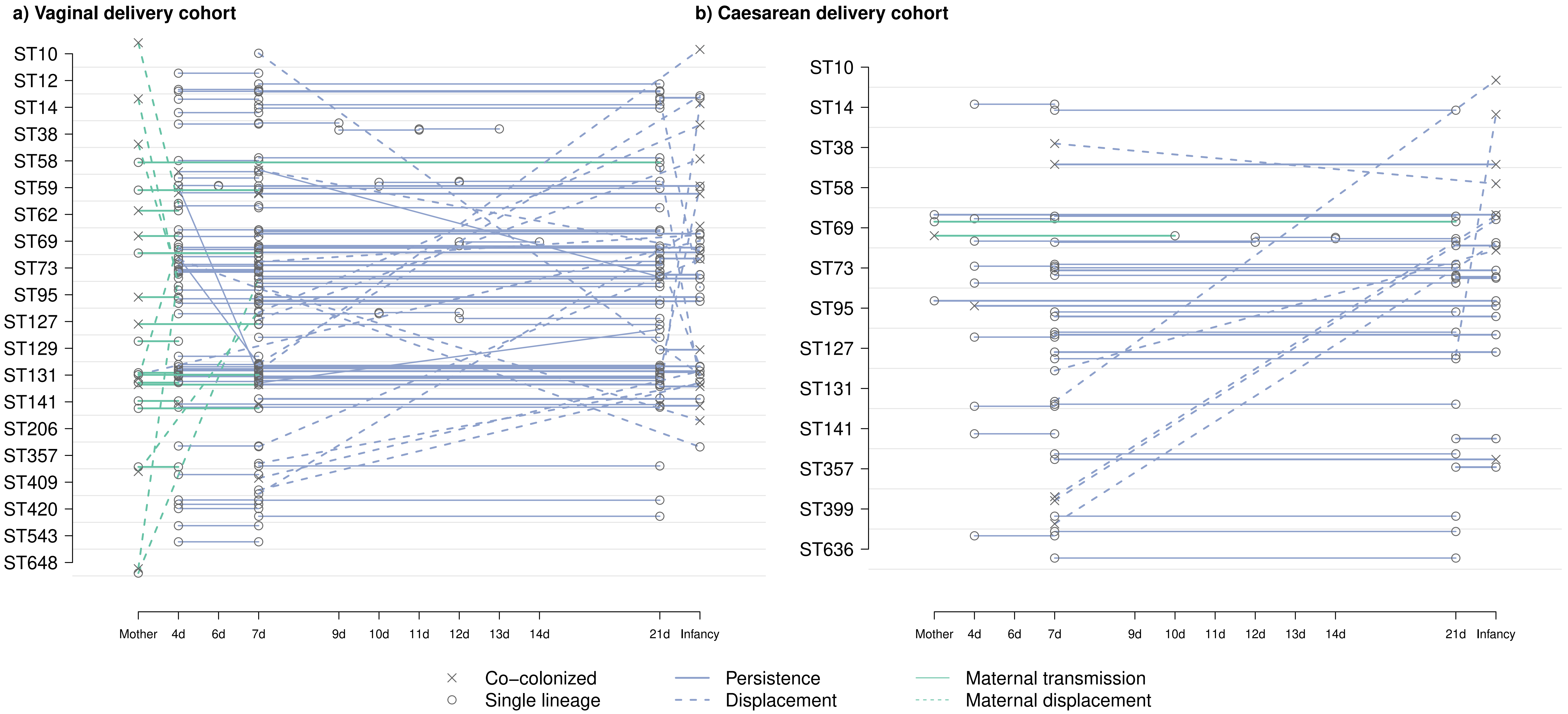


**Supplementary Figure 2 Longitudinal chart showing *Escherichia coli* lineage colonisation over time.** The plot shows positive identifications of *E. coli* sequence types (rows) in a sample taken at a certain time point (columns). Panel **a)** vaginal delivery cohort; panel **b)** caesarean section delivery cohort. Hollow circles represent reliable identifications of a single sequence type in the sample and black crosses identifications of coexisting sequence types. Connected solid or dashed lines represent the samples taken from a single individual (time points labelled with the number or days or 'Infancy') or their mother. A solid line connects samples where the same lineage was identified in two consecutive time points, and a dashed line connects samples where two different lineages were identified. Lineages that were identified in both the mother and a sample from the baby are connected by a solid light green line. Dashed light green lines connect samples, where the mother carried an E. coli lineage but the sample from the baby contained a different *E. coli* lineage. Horizontal solid lines signify identification of the lineage at several time points and angled dashed lines signify a switch from one lineage to another. Only lineages which were identified at least five times are shown.

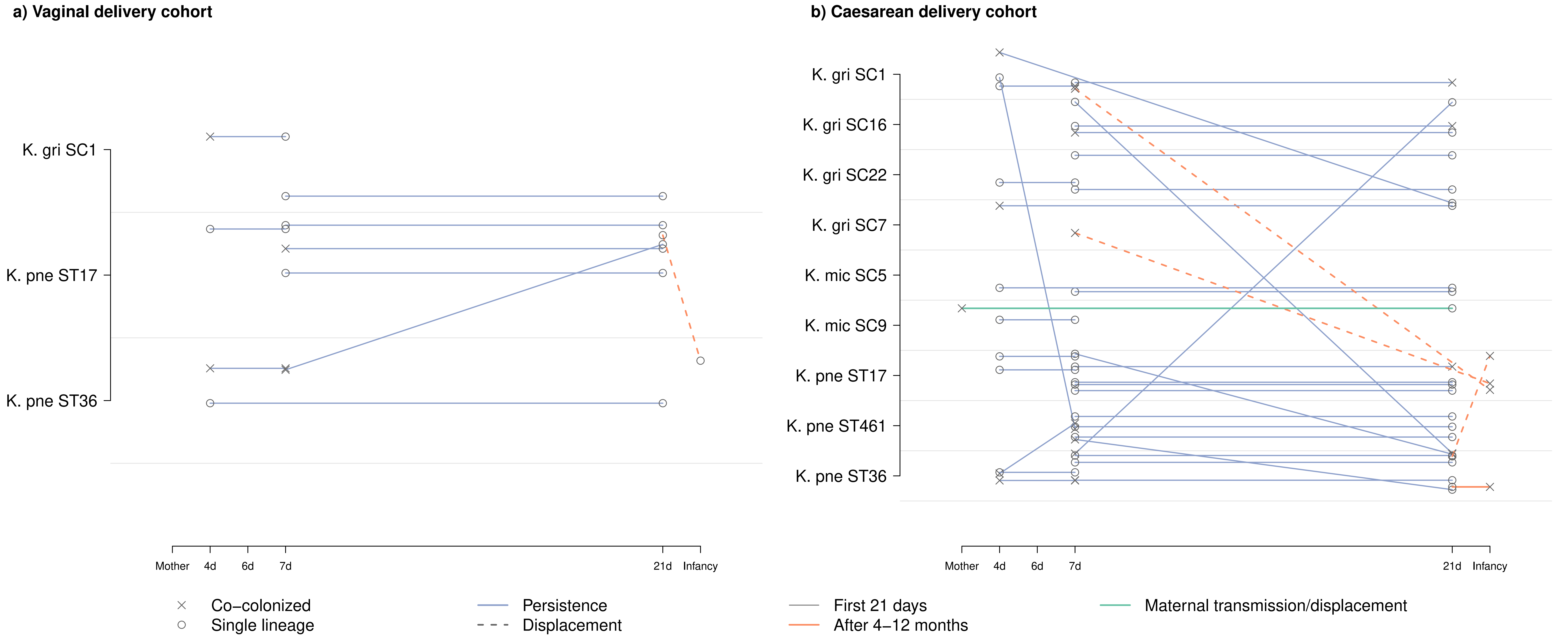


**Supplementary Figure 3 Longitudinal chart showing *Klebsiella* species and lineage colonisation over time.** The plot shows positive identifications of *Klebsiella* species and their sequence types (ST) or sequence clusters (SC) in the rows in a sample taken at a certain time point (columns). Panel **a)** vaginal delivery cohort; panel **b)** caesarean section delivery cohort. Hollow circles represent reliable identifications of a single sequence type in the sample and black crosses identifications of coexisting sequence types. Connected solid or dashed lines represent the samples taken from a single individual (time points labelled with the number or days or 'Infancy') or their mother. A solid light blue line denotes the samples taken within the first 21 days of life, a dashed light green line transmission/displacement from the mothers, and a dashed orange line the follow-up sampling done 4-12 months later. Horizontal lines signify identification of the lineage at several time points and angled lines signify a switch from one lineage to another. Only lineages which were identified at least five times are shown.

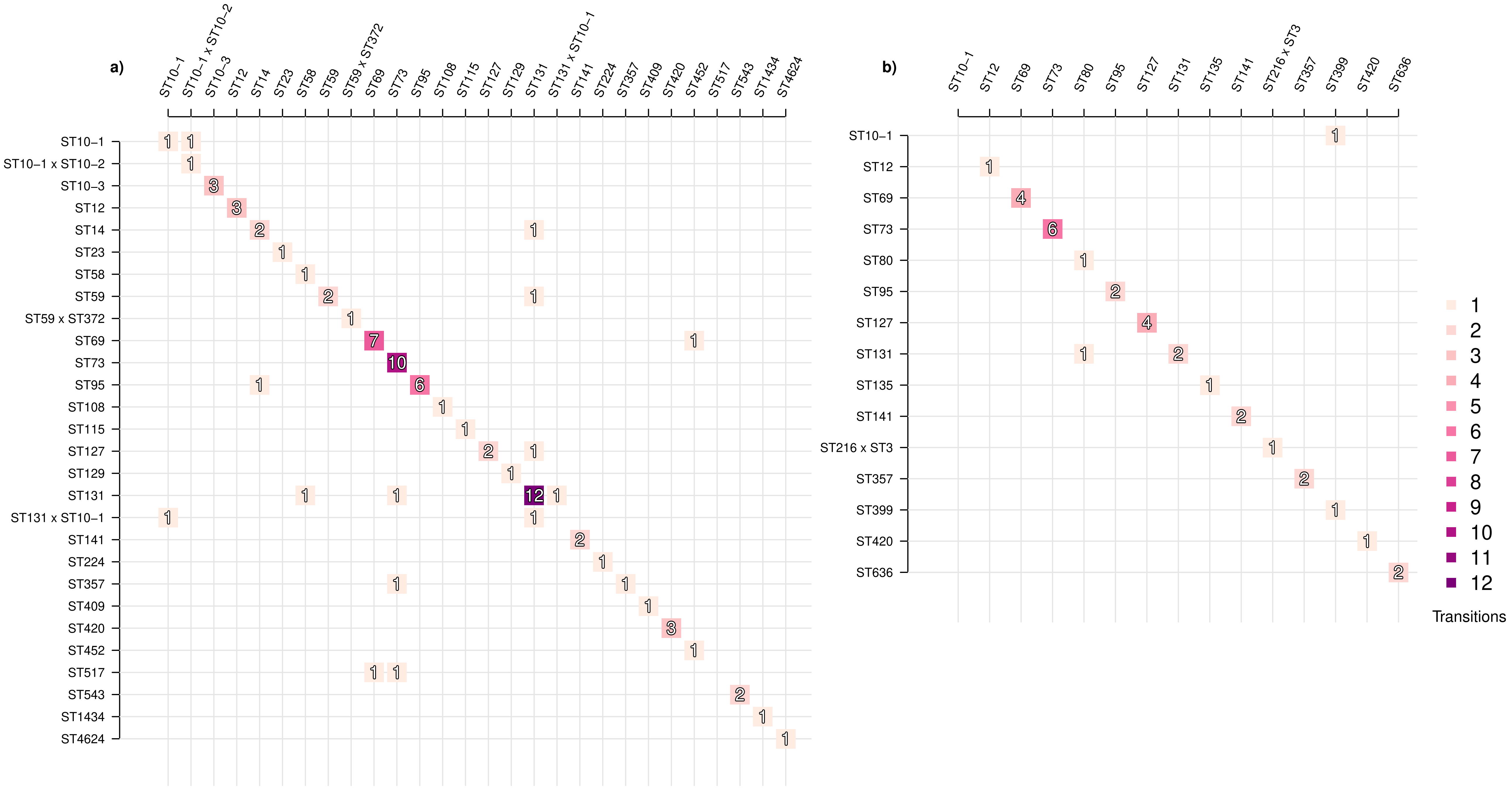


**Supplementary Figure 4 Event matrix displaying colonisation identities with respect to *Escherichia coli* lineages between subsequent time points.** The figure shows events corresponding to either transition from one *E. coli* lineage (rows) to another *E. coli* lineage (columns) or persistence of the same lineage (diagonal). Panel **a)** shows events for the vaginal delivery cohort with samples from the infancy period included, and panel **b)** shows the caesarean section delivery cohort with infancy period included. Darker shades of purple denote more common events, the count of which is also indicated by the number contained within the shaded boxes. Lineages shown were visited at least twice across the whole set of samples.

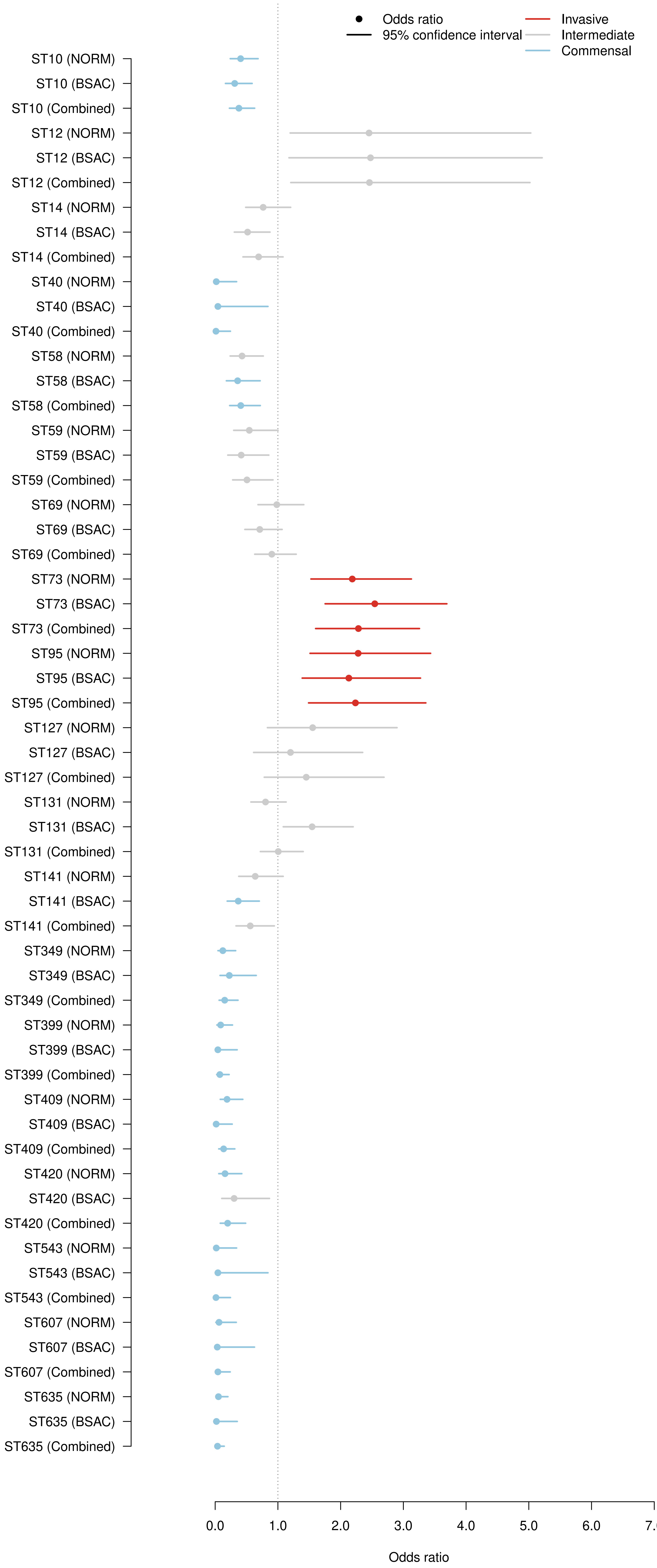


**Supplementary Figure 5 Full odds ratios for relative *Escherichia coli* invasiveness.** The odds ratios for invasiveness are displayed with the 95% confidence interval, where an OR of > 1 corresponds to more invasive and <1 to more commensal ST, for 19 lineages. The odds ratios are shown separately for either the NORM or the BSAC collection, or the combination of both. Lineages for which a significant OR was observed after correcting for multiple testing are coloured with a light blue or red colour.

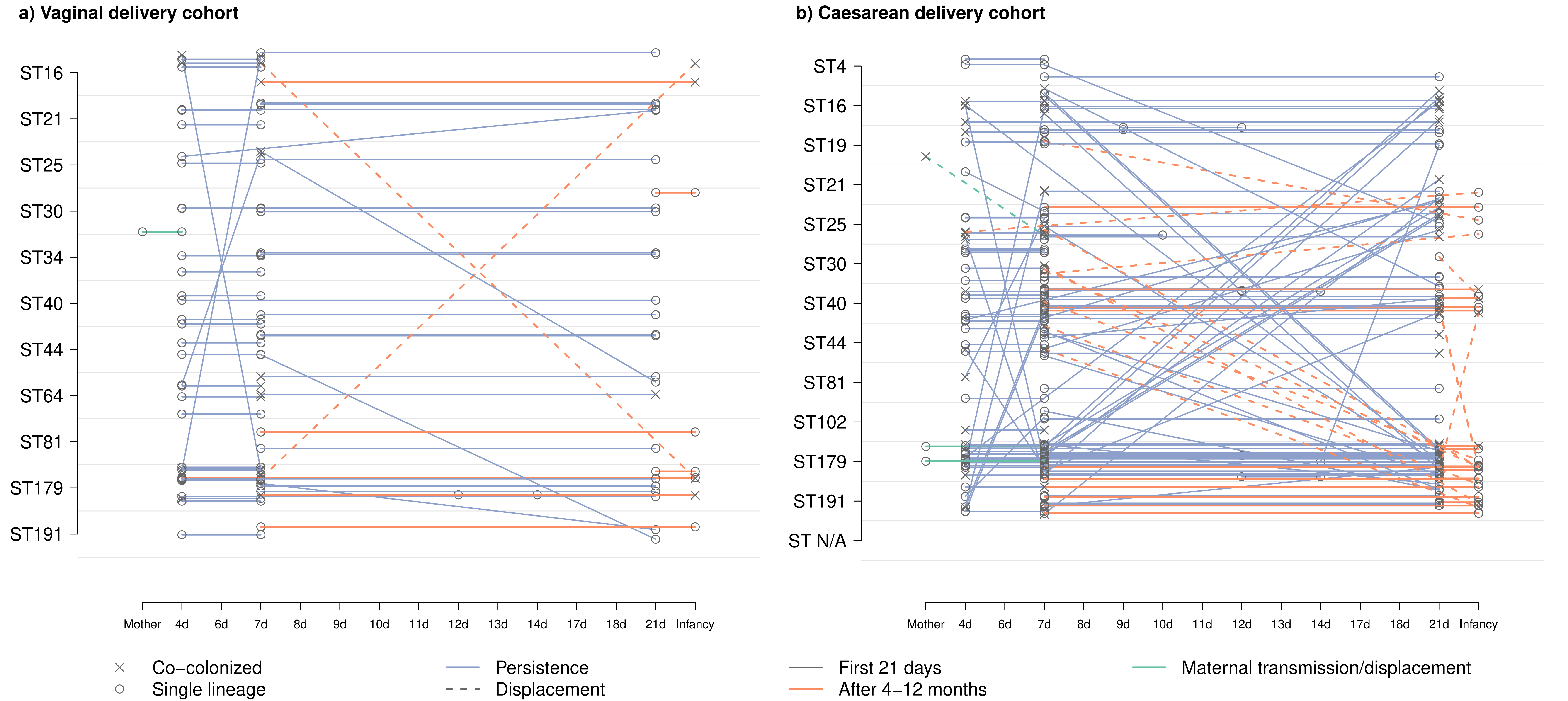


**Supplementary Figure 6 Longitudinal chart showing** ***Enterococcus faecalis*** **lineage colonisation over time.** The plot shows positive identifications of *E. faecalis* sequence types (rows) in a sample taken at a certain time point (columns). Panel **a)** vaginal delivery cohort; panel **b)** caesarean section delivery cohort. Hollow circles represent reliable identifications of a single sequence type in the sample and black crosses identifications of coexisting sequence types. Connected solid or dashed lines represent the samples taken from a single individual (time points labelled with the number or days or ‘Infancy’) or their mother. A solid light blue line denotes the samples taken within the first 21 days of life, a dashed light green line transmission/displacement from the mothers, and a dashed orange line the follow-up sampling done 4-12 months later. Horizontal lines signify identification of the lineage at several time points and angled lines signify a switch from one lineage to another. Only lineages which were identified at least five times are shown.

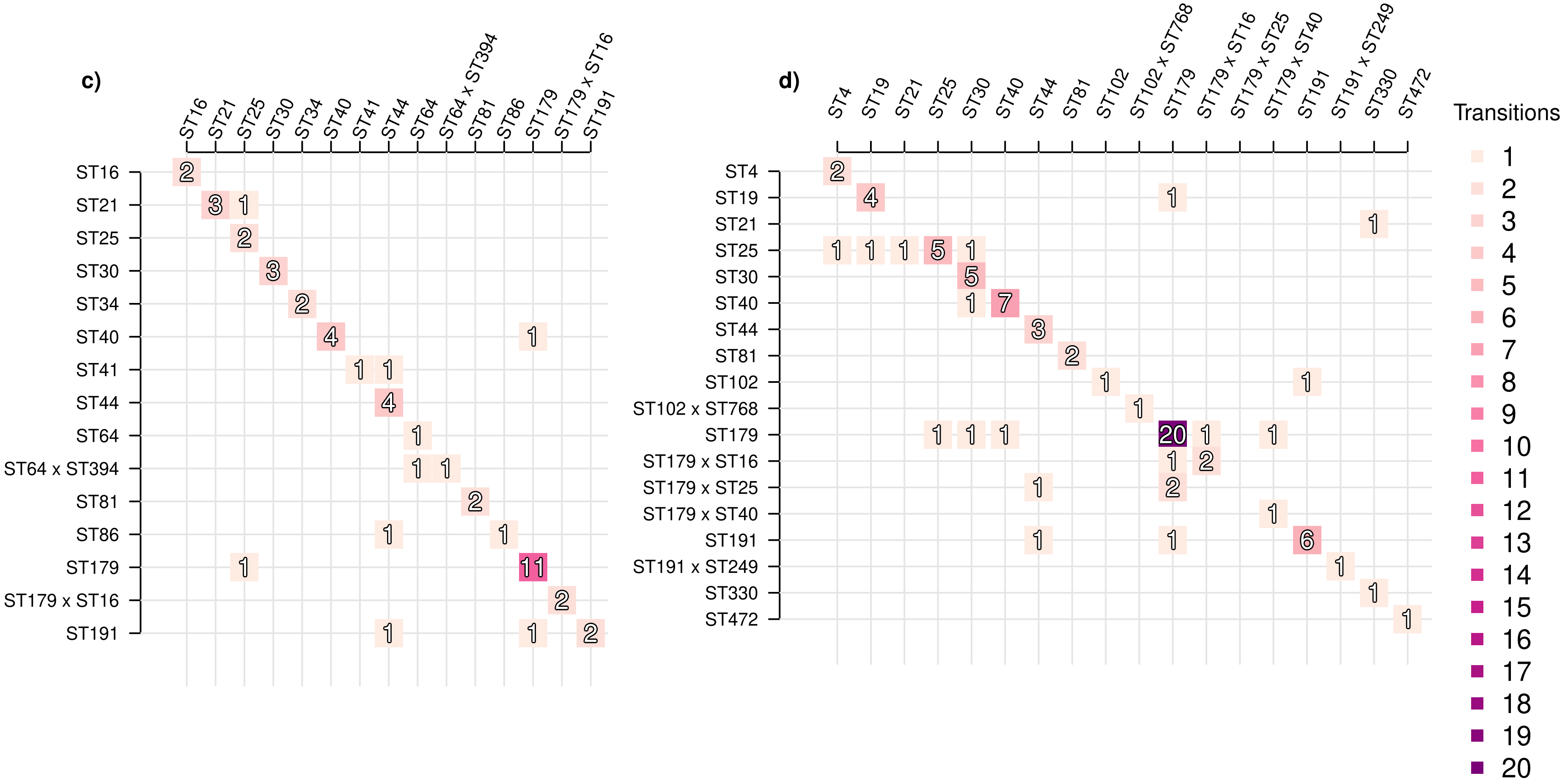


**Supplementary Figure 7 Event matrix displaying colonisation identities with respect to *Enterococcus faecalis* lineages between subsequent time points.** The figure shows events corresponding to either transition from one *E. faecalis* lineage (rows) to another *E. faecalis* lineage (columns) or persistence of the same lineage (diagonal). Panel **a)** shows events for the vaginal delivery cohort with samples from the infancy period included, and panel **b)** shows the caesarean section delivery cohort with infancy period included. Darker shades of purple denote more common events. Lineages shown were visited at least twice across the whole set of samples.
